# cohorts: A Python package for clinical ‘omics data management

**DOI:** 10.1101/626051

**Authors:** Nicholas P. Giangreco, Barry Fine, Nicholas P. Tatonetti

## Abstract

**Summary:** Precision medicine uses patient clinical and molecular characteristics to personalize diagnosis and treatment. This emerging discipline integrates multi-modal data into large-scale studies of human disease to make accurate individual-level predictions. The success of these studies will depend on the generalizability of the results, the ability of other researchers and clinicians to replicate studies, and the understandability of the methods used. Tools for data management and standardization are needed to promote flexible, transparent, and reproducible analyses. Here we present *cohorts,* a python package facilitating clinical and biomarker data management to enhance standardization and reproducibility of clinical findings.

**Availability:** The python package *cohorts* is available at http://www.github.com/ngiangre/cohorts.

## 1 Introduction

Precision medicine improves patient diagnoses and treatments by integrating clinical and biomarker data, which requires effective data management for reproducibility of clinical research findings (Vargas and Harris, 2016; Leopold and Loscalzo, 2018; Niven *et al.*, 2018). The studies often span institutions, departments, and research teams, where patient data is collected and processed in a specialized way. Clinical studies including patient data from multiple sources are integrated and managed by the primary research team, and so correct attribution of the clinical and biomarker characteristics is critical. Feasible and accessible software can facilitate data storage and management, which promotes flexible, transparent, and reproducible analyses. Moreover, software that facilitates integration of multiple patient cohorts can promote the use of advanced statistical and machine learning analyses.

Precision medicine calls for rigor and reproducibility in data generation and data analysis (Mueller *et al.*, 2018; Orton and Doucette, 2013). For large consortium based studies, computational platforms allow for sharing data and merging of multisite datasets (Lam *et al.*, 2016). While large-scale studies may have an extensive computing platform with custom software (Wolstencroft *et al.*, 2015; Price *et al.*, 2019), most clinical studies are smaller and would benefit from standardized procedures and interoperable data management and analysis tools. There are many software packages and platforms that perform precision medicine data analysis, such as pyGeno (Daouda *et al.*, 2016) and in particular packages from the package manager Bioconductor in the R programming language (Huber *et al.*, 2015). However, many of these applications can handle only one experimental type or are particular to a specific end-to-end data analysis. There is a need for flexible and open source software, where patient clinical and biomarker data can originate from different cohorts, platforms, and data types.

Here, we present the python package *cohorts,* which provides storage and management of patient clinical and biomarker data. Moreover, *cohorts* facilitates integration of multiple patient cohort data through common attributes shared by each cohort. *Cohorts* is light-weight and portable software promoting standardized and reproducible clinical research analyses. We provide this open source python package on Github at http://www.github.com/ngiangre/cohorts.

## 2. Results

### 2.1 Package overview

The main objective of *cohorts* is for the storage and management of patient clinical and biomarker datasets (Fig. 1). The management of patient data is through the python class **Cohort** (classes are denoted in bold). **Cohort** takes as input file locations of 1) a replicate or sample dataframe, where rows are biomarkers and columns are replicates or samples (names of rows and columns are marker ids and replicate or sample names, respectively), and 2) a sample groups dataframe, where rows are the clinical outcomes or groups (e.g. reference and treatment; can be any data type) and columns are the samples (names of rows and columns are the group names and the sample names, respectively). In addition to the name of the cohort and additional parameters (see package for details), **Cohort** derives and stores attributes of the dataset in an object-oriented fashion, containing both common attributes to other declared classes of **Cohort** as well as specific attributes for the patient cohort to be used in downstream analysis. The data management by the patient cohort object allows for standardized analysis with minimal initial data upload and wrangling. Moreover, the python object gives consistent and logical data attributes that are easily retrievable for downstream analyses.

**Figure 1.**
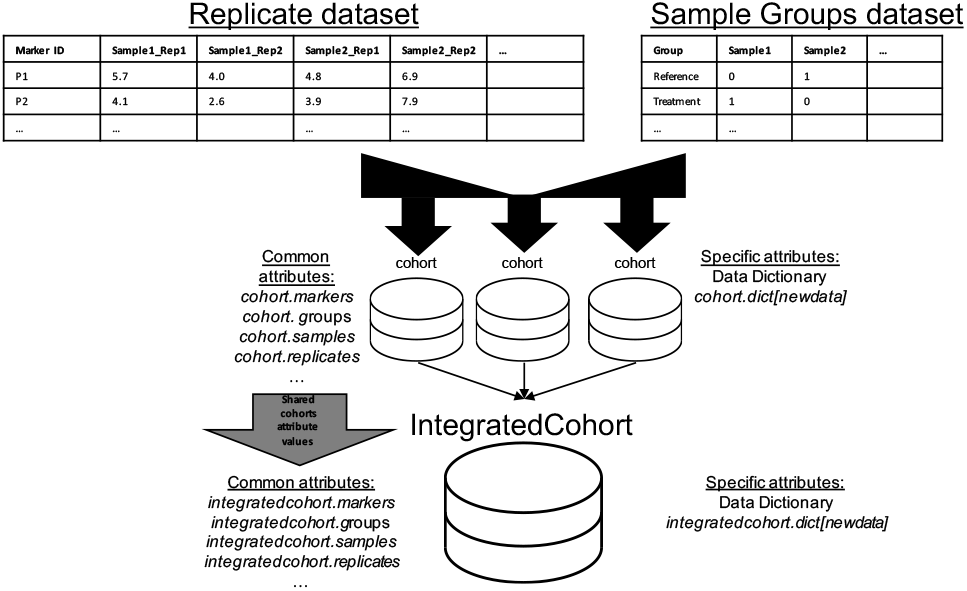
Schematic of data input, storage, management, and integration workflow for the *cohorts* package.

Integration of multiple datasets is through the **IntegratedCohort** python class. By providing a dictionary of **Cohort** objects, **IntegratedCohort** parses, integrates, and containerizes shared information from each **Cohort** object. This class declaration is created by using the common data attributes of the **Cohort** class (at least one **Cohort** object is needed to produce an **IntegratedCohort** object). Additionally, specific attributes are preserved for the individual patient cohorts and are available for the newly created **IntegratedCohorts** object. Specific attributes for this object can facilitate, for example, managing data from statistical and machine learning tasks using shared biomarkers or clinical groups. Again, no initial data upload and minimal wrangling is needed for this integration task, saving the user time to focus on standardized and reproducible downstream biomarker analysis.

### 2.2 Implementation

Implementation of the *cohorts* package is described and shown in the supplementary material in the online version of the manuscript. Briefly, data for an individual patient cohort is managed through the **Cohorts** class. The patient data is retrieved through accessing object attributes, and new data can be stored within the data attribute. The **IntegratedCohorts** class integrates patient data stored in *Cohorts* objects using the “integrate_cohorts” method, which canaccess, parse, integrate, and store all the patient data within the **IntegratedCohort** class. Example procedures streamlined using the package include descriptive statistics of molecular marker data and outcome prediction using integrated patient data that share common attributes. An example can be found in a jupyter notebook at https://github.com/ngiangre/cohorts/blob/master/Bioinformatics_Note_Implementation.ipynb.

## 3. Conclusion

Studies in precision medicine need standardized and reproducible analyses. The *cohorts* package supports data management, procedural standardization, and data integration for analyses that will improve the flexibility, transparency and reproducibility of clinical investigations. We provide jupyter notebooks giving an overview and usage of the package in the *cohorts* GitHub repository at http://www.github.com/ngiangre/cohorts.

## Acknowledgements

Thank you to the Columbia University Irving Medical Center Proteomics Core for generation and pre-processing of patient proteomic samples. Most importantly, thank you to the patients for consenting to partaking in clinical research.

## Supplementary Material

### Implementation

Implementation of the *cohorts* package is described and shown below. The source code can be found at https://github.com/ngiangre/cohorts/blob/master/Bioinformatics_Note_Implementation.ipynb.

### Patient cohort data

The **Cohorts** class manages data from a single patient cohort. The package is accompanied by a synthetically generated patient cohort dataset at https://github.com/ngiangre/cohorts/tree/master/sample_data. The data, which in this example will include protein biomarker expression, follows the necessary format outlined in section 2.1. After importing the *cohorts* package, the patient cohort data is uploaded by specifying directory and file locations as values and the cohort name as the key within a dictionary. The dictionary is then given as a parameter to the **Cohort** class constructor to instantiate a patient cohort data object. Within the **Cohort** class, the patient cohort data is automatically parsed and stored as attributes in the newly created object. The patient data can then be retrieved through accessing object attributes, and new data can be stored within the data attribute. For example, the distribution of protein expression across patients can be accessed and displayed (Fig. 2).

**Figure 2.**
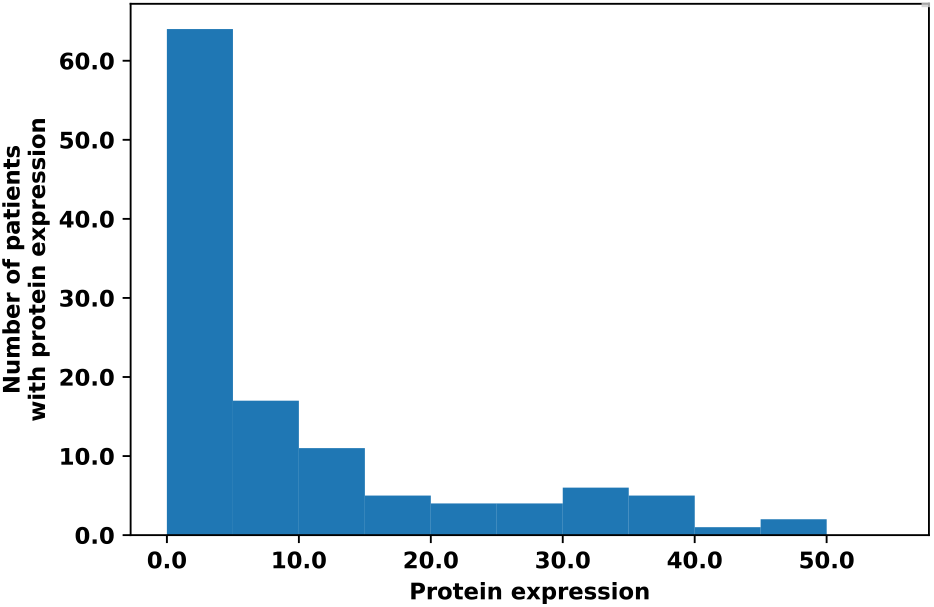
Protein expression distribution across patient cohort samples using the *cohort* package. See Implementation jupyter notebook for source code.

### Integration of multiple patient cohort data

The **IntegratedCohorts** class integrates patient data stored in *Cohorts* objects. As shown in the notebook, a dictionary of three different patient cohort datasets is input to create an instance of the **IntegratedCohorts** class. After calling the ‘integrate_cohorts’ method the original patient cohort data is accessed, parsed, integrated, and stored within attributes of the **IntegratedCohort** class. In addition to data integration, the individual patient cohort data is stored separately upon instantiation. The integration of related patient biomarker and clinical data provides the ability to generalize findings from an individual to a broader patient population. An example is the task of disease prediction. The characterization of disease by biomarkers or clinical markers may only be representative in a patient population, but providing similar results within a larger patient population provides stronger evidence for association of patient markers to a disease. Figure 3 shows a receiver operator characteristic curve from predicting disease from protein marker expression levels of patient samples in the three patient cohort datasets.

**Figure 3.**
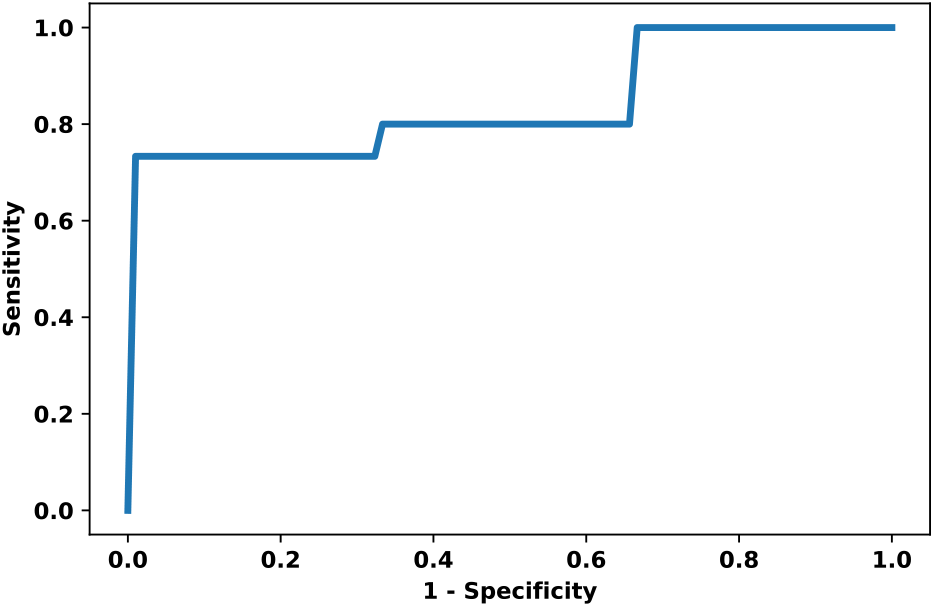
Receiver-operator characteristic curve for disease prediction using protein marker expression as features in a Logistic Regression classifier. The average sensitivity is shown from 5-fold cross validation.

